# Hearing what is being said: The distributed neural substrate for early speech interpretation

**DOI:** 10.1101/2023.03.23.533971

**Authors:** Alex Clarke, Lorraine K. Tyler, William Marslen-Wilson

## Abstract

Speech comprehension is remarkable for the immediacy with which the listener hears what is being said. Here, we focus on the neural underpinnings of this process in isolated spoken words. We analysed source-localised MEG data for nouns using Representational Similarity Analysis to probe the spatiotemporal coordinates of phonology, lexical form, and the semantics of emerging word candidates. Phonological model fit was detectable within 40-50 ms, engaging a bilateral network including superior and middle temporal cortex and extending into anterior temporal and inferior parietal regions. Lexical form emerged within 60-70 ms, and model fit to semantics from 100-110 ms. Strikingly, the majority of vertices in a central core showed model fit to all three dimensions, consistent with a distributed neural substrate for early speech analysis. The early interpretation of speech seems to be conducted in a unified integrative representational space, in conflict with conventional views of a linguistically stratified representational hierarchy.

## Introduction

Human speech comprehension is remarkable for the earliness with which a lexically and semantically interpreted speech percept becomes available to the listener, at lags of 250 ms or less (Marslen-Wilson, 1973, 1975, 1987; Marslen-Wilson & Tyler, 1975). Equally remarkably, despite 50 years of research, two foundational questions remain in dispute: What is the basic functional and neurobiological architecture that underpins these early interpretative processes and what is the computational vocabulary in terms of which the human brain conducts these operations?

The currently and historically dominant view in the cognitive neurosciences takes for granted that a *linguistically stratified representational hierarchy,* centred round the concept of the phoneme, mediates the link between sound and meaning. On this account, acoustic information in the speech stream is mapped, most likely through processes of featural decomposition (e.g. Mesgarani et al., 2014; Warren & Marslen-Wilson, 1987), onto a distinct pre-lexical level of phonological representation specified in phonemic terms. The resulting sequence of phoneme labels provides access to stored representations of lexical form, which link in turn to the meaning representations associated with these forms. The neurobiological realisations of this approach distinguish an anatomically and computationally distinct hierarchy of primarily feedforward processes, radiating posteriorly and anteriorly from spectrotemporal analyses in Heschl’s gyrus to phonetic feature extraction in the superior temporal gyrus (STG), leading to phoneme identification and thence to potential lexical forms and meanings in the middle temporal gyrus (MTG) (e.g. DeWitt & Rauschecker, 2012; Di Liberto et al., 2015; Gwilliams et al., 2018; Gwilliams & Davis, 2022; Hickok & Poeppel, 2007; Leonard & Chang, 2014).

This stratified separation of processes operating on the acoustic-phonetic input and those operating on representations of lexical form and meaning is inconsistent, however, with longstanding behavioural evidence (e.g. Gaskell & Marslen-Wilson, 1996, 2002; Marslen-Wilson & Warren, 1994), as well as more recent electrophysiological evidence (e.g. Cibelli et al., 2015; Leonard et al., 2015; Yi et al., 2019), that the basic phonological interpretation of the speech input is intrinsically modulated by the lexical and sentential environments in which these sounds occur and in which the listener has learned, from infancy, to discriminate them. The stratified approach fails, furthermore, to resolve - or even consider - the further critical role of speech information in constructing the listener’s perceptual representation of speech as they hear it, assuming instead that phonological analyses fade away once they have fulfilled their role in mediating access to meaning.

A contrasting approach, which meets both these concerns, was pioneered by the Distributed Cohort concept (Gaskell & Marslen-Wilson, 1999, 1997, 2002), based on a *fully distributed neural architecture*, where acoustic-phonetic information was mapped directly, via a recurrent Elman net, onto parallel output representations of lexical semantics and lexical phonology. This implementation required the connection weights in the network to code information about both mappings such that the process of identifying a word involved the early and continuous integration of both types of constraint. Yi et al (2019) make a related, neurobiologically more explicit set of proposals, also incorporating Elman recurrent nets, but focus primarily on the computation of a contextually modulated level of phonological representation in STG, with less emphasis on broader processes of speech interpretation (also see Hamilton et al., 2021). The Distributed Cohort (DisCo) approach that we develop here does not assume a comparable neurocomputationally distinct phonological analysis of the speech input, but rather a representationally more neutral process that converges onto an integrated phonological and lexicosemantic output as acoustic-phonetic constraints propagate through the network - and where this output forms the basis for the listeners’ perceptual interpretation of the speech they hear.

These two kinds of approach make very different claims about the overall functional architecture of this process, in terms of the nature, the timing and the relative locations of the different processes underpinning the early stages of speech interpretation. The failure, nonetheless, of decades of research to resolve such basic disagreements about how the brain relates sound to meaning, reflects the historical absence of time-resolved experimental tools capable of mapping the actual spatiotemporal distribution of neural processes engaged as speech is heard, and of determining the computational content of these processes (Marslen-Wilson, 2019).

More recent research, using either non-invasive methods based on source localised MEG signals or more invasive methods (ECOG) recording directly from the brains of surgical patients, has not yet provided a unified bihemispheric picture that meets these requirements. An early study by Travis et al. (2013), for example, combined evidence from MEG and ECOG to show rapid processing of phonetic speech sounds from 60 ms after word onset in superior temporal cortex and much later semantic processing of the spoken word after it had been uniquely identified. The study did not, however, address the mechanisms whereby early phonetic analyses link to processes of cohort access and competition. Subsequent MEG research, combining source-localised MEG data with multivariate Representational Similarity Analysis (Kocagoncu et al., 2017), focused on the dynamic processes of lexicosemantic analysis prior to accessing the meaning of the target word, but did not examine the early neurocomputational transition, aligned to word-onset, whereby the initial speech input is related to these key lexical and semantic processes. A different approach, combining MEG data from continuous speech with linear kernel estimation to detect processing effects of relevant cognitive variables (Brodbeck, Presacco, et al., 2018), shows early sensitivity to acoustic variables at around 70 ms from word onset, followed by effects of phoneme surprisal and cohort entropy at 110 and 120 ms in superior temporal cortex (Brodbeck, Hong, et al., 2018). But because these (and related) studies compute the onset of phonological and lexical activation for words heard in extended discourse contexts, they cannot separate the spatiotemporal signature of bottom-up stimulus-driven early analyses from the potential effects of the contexts in which these words are heard.

These primarily MEG-based studies are complemented by the remarkably detailed but punctate picture of lower-level speech analysis processes, viewed as a mosaic of multiple specialized processors, that is generated by ECOG methods (Leonard & Chang, 2014; Yi et al., 2019). Mesgarani et al., (2014), for example, showed that the identification of different phonetic features was supported by distinct electrode sites, but where these sites displayed no feature-specific spatial clustering, reflecting an intermixed mosaic of distinct processing areas. Yi et al. (2019) argue that these anatomically scattered STG populations encode multiple sources of information which are dynamically integrated over time. Hamilton et al (2021) provide extensive evidence against a conventional serial hierarchy linking primary auditory cortex to linguistically relevant analyses in lateral STG, while also arguing for a distributed mosaic interpreting different acoustic and phonetic cues in the speech signal. However, as noted above, these studies generally assume a stratified linguistic level of representation, characterized as phonological in nature, and have relatively less to say about how this level relates to neural representations of lexical form and meaning.

To tap more directly into the broader neural framework underpinning the early stages of speech interpretation, the current study uses spatiotemporal searchlight Representational Similarity Analysis (ssRSA; Kocagoncu et al., 2017; Lyu et al., 2019; Su et al., 2012, 2014) in MEG source space to track the spatiotemporal coordinates, for words heard in isolation, of the neurocognitive processes that underpin the activation of the phonological, lexical and semantic properties of the word-initial cohort. The choice here of a research strategy based on MEG source space and searchlight-based RSA is consistent with the ‘Hopfieldian’ assumption that complex cognitive functions can only be realised by the distributed population structure of neural state spaces and the system’s movement within and between these spaces (Barack & Krakauer, 2021; Dubreuil et al., 2022; Saxena & Cunningham, 2019).

The singular advantage of ssRSA in this context is that it allows direct comparison of patterns of neural activity across the brain with the predictions of different theoretical models of the neurocomputational content of these processes. In doing so it assumes (a) that the ‘representational geometry’ of neuronal population codes captures the structured pattern of neural activity whereby the brain represents the properties of a particular type of content, and (b) that the RSA method of representational dissimilarity matrices (RDMs) captures, in an abstract format, the distinguishing properties that define the representational geometries of the different dimensions at issue (Kriegeskorte et al., 2008; Kriegeskorte & Diedrichsen, 2019; Kriegeskorte & Kievit, 2013).

This means that the spatiotemporal pattern of RSA model fit, for a given model RDM over a given area of searchlight analysis, can tell us not only *where* and *when* we see relevant patterns of neural activity but also *what* is the likely neurocomputational content of the processes associated with these areas of model fit. Here we exploit this to capture the spatiotemporal coordinates of the early phonological analysis of the speech input and to determine how this analysis relates to the timing and location of access to the potential lexical identities (the word-initial cohort) signalled by these accumulating phonological cues and to the semantic properties associated with these early cohort members.

To do this requires discrete quantifiable measures of each of these different aspects of form and meaning, which we can then separately test, using ssRSA, against the temporally evolving neural activity as a word is heard. To capture the phonological properties of the speech input we utilise the articulatory phonetic features of speech sounds. There is converging evidence across imaging modalities that major subgroups of neurons in superior temporal cortex are sensitive to articulatory phonetic features (Correia et al., 2015; Di Liberto et al., 2015; Hamilton et al., 2021; Mesgarani et al., 2014; Obleser et al., 2004; Tang et al., 2017; Wingfield et al., 2017), and that such features link sensory signals to more abstract lexicosemantic representations (Hickok & Poeppel, 2007; Leonard & Chang, 2014). Because we know what speech sounds the listeners are hearing, a phonological model capturing the properties of this stimulus ensemble can be computed directly from a specification of these sounds in terms of their articulatory features (see Figure 1A). In doing so, however, we do not commit to the processing role of these features as abstract units of neural computation, but rather as a way of capturing the sequential articulatory patterns underlying the production of the speech waveform.

**Figure 1.**
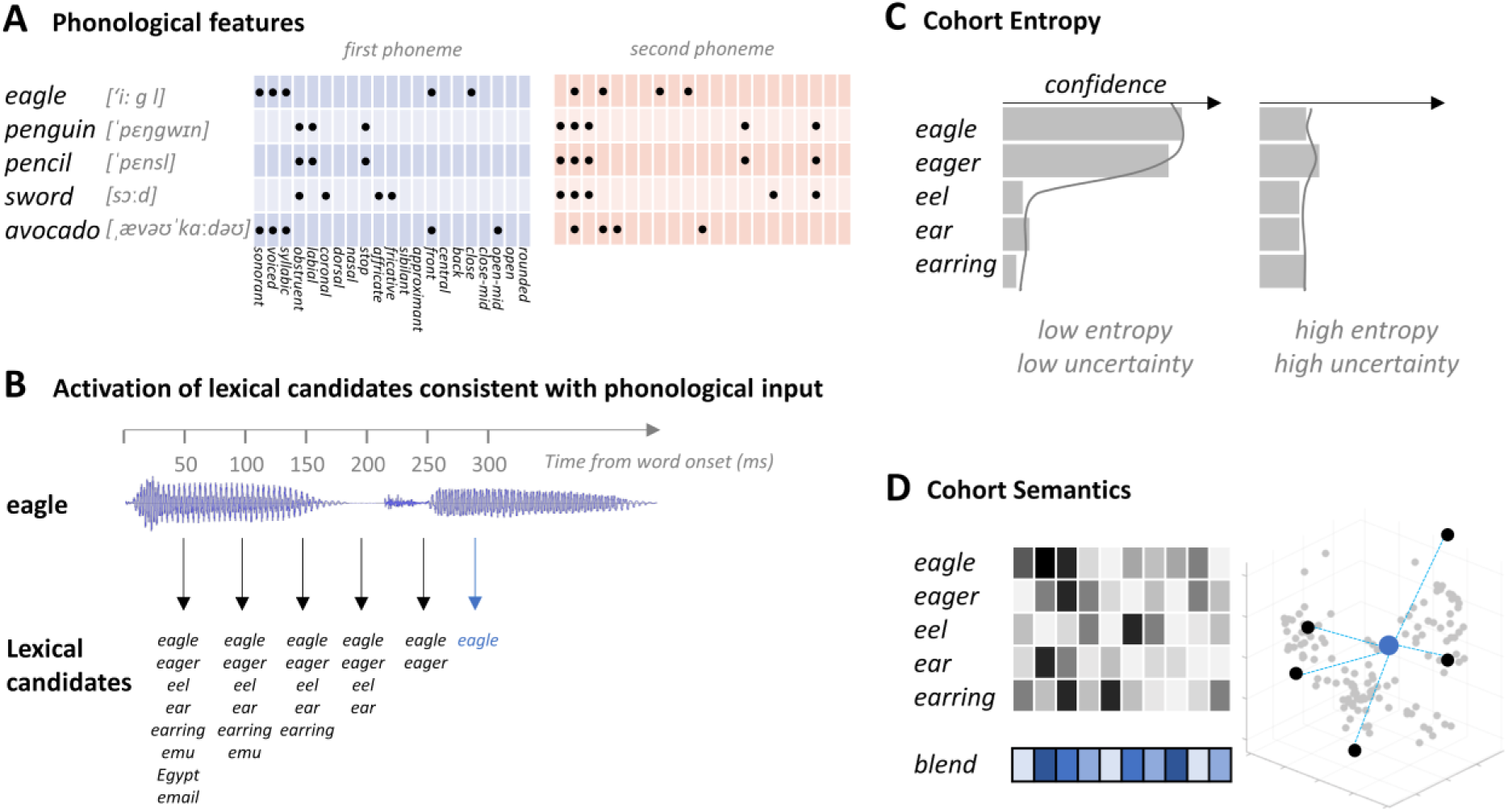
Cognitive measures and models. A) Phonological articulatory features are derived for the first and second phoneme of each word, specifying which of the 21 articulatory features apply to each sound. B) The gating procedure used to define the candidates based on increasing segments of a spoken word, illustrated with the example of ‘eagle’. Early gates (e.g. 50 ms) generate several candidates beginning with the same onset sound and the candidate set decreases as more speech is heard. The point in time when a single candidate remains is considered the recognition point – indicated by the blue arrow. C) Cohort entropy is calculated from the confidence values across candidates at a given gate, resulting in low entropy when participants are more certain of the word identity and high entropy when they are less certain. D) A blended measure of the semantics of cohort members was calculated from a corpus-based distributional semantic vector for each candidate. The blended semantic vector is created by averaging the vectors of the candidate set for each word fragment.

To separate analytically the lexical activation of a word-initial cohort from the activation of the semantics of that cohort, prior to unique word recognition, we developed two further models tapping selectively into these two aspects of the recognition process (Figure 1). This required an estimate of the membership of the word-initial cohort, hypothesised to be covertly generated as the initial phonetic input is heard. To achieve this, a behavioural gating task was used where incremental word fragments were heard and participants were asked to say which word they thought they were hearing at successive 50 ms increments (as in Klimovich-Gray et al., 2019).

Based on the resulting word-initial cohorts, evolving across gates for each word in the study (see Figure 1B), we first calculated a Cohort Entropy measure reflecting the lexical uncertainty of word identity based on the speech input for that gate (Figure 1C). A high entropy initial speech sound, consistent with many possible words, generates high uncertainty, and conversely for low entropy initial sounds. This is a measure that cannot be computed based on phonological information alone, since it depends on the potential lexical implications of this information. At the same time, it does not depend on the semantic properties of candidate words, since the critical variable is the number and likelihood of candidates generated at each gate for each word and not the meaning of these candidates.

In contrast, to detect the activation of semantic information associated with early word candidates, estimates of the semantic content of each candidate were combined to provide a single summary semantic representation of the entire candidate set for each word at each gate (Cohort Semantics; Figure 1D). To achieve this, a distributional semantic representation of each candidate was obtained from corpus-based computational analyses (Baroni & Lenci, 2010) and combined across the cohort, resulting in a blended measure of cohort semantics (Gaskell & Marslen-Wilson, 1999; Klimovich-Gray et al., 2019). Any relationship between neural activity and this semantic measure will indicate that semantic information related to the current cohort members is in the process of being accessed.

In summary, the goal of this study is to characterise the properties of strictly bottom-up processes of early speech analysis, independent of potential constraints or predictions from a prior context. Using ssRSA analyses of the MEG signals based on the three models described above, we aim to provide a differentiated spatiotemporal map for three critical dimensions underlying early speech interpretation - the phonological interpretation of acoustic-phonetic information in the speech signal, the implications of this information for potential lexical identities intended by the speaker, and the early access of semantic information related to these initial cohort members. The degree of spatial and temporal overlap between these three maps will help assess the viability of a stratified representational model as opposed to a fully distributed and representationally more neutral account of these core interpretative processes.

## Methods

The analyses reported in this paper were conducted on previously collected data from Kocagoncu, Clarke, Devereux & Tyler (2017), which focused on the end-stages of the word recognition process (the ‘recognition point’) for spoken words heard in isolation. Detailed methods are provided in the source paper, with the essential aspects repeated here.

### Gating Experiment

To define the word candidates initially elicited by the onsets of each target word, we conducted a behavioural gating study (Grosjean, 1980; Tyler & Wessels, 1985). In a self-paced procedure, 14 participants, who did not take part in the MEG study, listened to spoken fragments of the words used in the MEG study and were asked to type in their best guess of the word and rate their confidence in their answer on a scale of 1 to 7 (7: very confident; 1: not confident at all). Words were presented at incrementally increasing fragment lengths (i.e., gates) of 50 ms, 100 ms and 150 ms from vocal onset. The participants’ responses were used to obtain a list of candidates for each target word (Figure 1B), with a median of 9, 8 and 6 candidates per target word for the three fragment lengths (mode; 10, 8, 6; range; 2-14, 1-13, 1-12 respectively). The gating data were used to calculate measures of cohort entropy and cohort semantics (see *Cognitive Measures* below).

### MEG Participants, stimuli, and procedure

Eleven healthy participants (mean age 24.4 years, range 19-35 years, 7 females) took part in the main study. All were right-handed, native British English speakers with normal hearing. The experiment was approved by the Cambridge Psychology Research Ethics Committee and all participants gave informed consent.

The stimuli consisted of 218 spoken words in isolation, each of which was a concrete concept (e.g. *alligator, hammer, cabbage*), and 30 phonotactically legal nonwords (e.g. *rayber, chickle, tomula*). All words were spoken in a female voice, and all nonwords were excluded from the imaging analysis. Participants were instructed to listen to the speech stimuli and press a response key whenever they heard a nonword (10% of trials). Each trial began with a fixation cross for 650 ms, before the auditory stimulus, which was followed by a random inter-stimulus interval lasting between 1500 and 2500 ms. Blink periods were included at the end of each trial, lasting 1500 ms, during which an image of an eye appeared in the middle of the screen. Stimuli were presented using E-Prime 2 (Psychology Software Tools) and delivered binaurally through MEG-compatible ER3A insert earphones (Etymotic Research Inc., IL, USA). The delay in sound delivery due to the length of earphone tubes and the stimulus delivery computer sound card was 32 ms. This was corrected for during MEG preprocessing. Stimuli were presented in two blocks, each containing 109 words and 15 nonwords and were presented in a pseudo-randomized order. The block order was counterbalanced across participants.

### MEG and MRI acquisition

Continuous MEG signals were recorded using the whole-head 306-channel Vector-view system (Elekta Neuromag, Helsinki) with 102 planar gradiometer pairs and 102 magnetometers. Electro-oculogram (EOG) and electrocardiogram (ECG) electrodes were used to record eye blinks and cardiac activity, and five head position indicator (HPI) coils recorded the position of the head within the MEG helmet every 200 ms. The participants’ head shape was digitally recorded using a 3D digitizer (Fastrak Polhemus Inc., Colchester, VA), along with the positions of the EOG electrodes, HPI coils, and fiducial points (nasion, left and right periaricular). MEG signals were recorded continuously at 1000 Hz sampling rate with a high-pass filter of 0.03 Hz. To enhance source localization, T1-weighted MP-RAGE scans with 1 mm isotropic resolution were acquired for each subject using Siemens 3-T Tim Trio. Both the MEG and MRI systems were located at MRC Cognition and Brain Sciences Unit in Cambridge, UK.

### MEG preprocessing and source localization

Raw MEG signals were processed using MaxFilter 2.2 (Elekta Oy, Helsinki, Finland) to apply temporal signal space separation and head motion correction. Independent components analysis using EEGLAB was used to remove blink and cardio-correlated signals (Delorme & Makeig, 2004). Data were further processed using SPM8 (Wellcome Trust Centre for Neuroimaging, University College London, UK). Data were low-pass filtered at 40 Hz using a 5th order Butterworth filter, high-pass filtered at 0.5 Hz using a 5^th^ order Butterworth filter, downsampled to 250 Hz, epoched between −200 and 500 ms relative to word onset, and baseline corrected using the pre-stimulus time period.

Source localisation of MEG signals used a minimum-norm procedure based on both magnetometer and gradiometer sensors in SPM12. First the individual’s MRI image was segmented and normalised to the MNI template brain, before the canonical cortical head meshes were inverse normalised to the individual subjects’ head space. The cortical mesh consisted of 5124 vertices corresponding to dipoles with fixed orientations perpendicular to the cortical surface. MEG sensor locations were co-registered to the MRI image using the three fiducial points (nasion, left preauricular, right preauricular) and the additional headpoints recorded prior to MEG. A single shell forward model was used, and the data from both magnetometers and gradiometers were inverted together (Henson et al., 2009) using a minimum-norm estimate (IID in SPM) and the default settings (with the exception that no hamming window was applied) resulting in an activation time-course at each vertex on the cortex.

### Representational similarity analysis (RSA)

RSA was used to compare the predicted similarity between the word stimuli, based on our cognitive measures, to the similarity derived from MEG patterns. ssRSA (Su et al., 2012) provides a way in which the predictions embodied by the representational geometry of our cognitive measures can be tested against the representational geometry of time-evolving neural activity patterns.

### Cognitive measures

In this study, we are interested in the initial responses to spoken words, prior to the word recognition point. To this end, we modelled the phonological features of each word, in addition to the entropy and semantics of the lexical word candidates of an initial cohort (Figure 1; see gating task).

The *phonological feature model* was based on a phonetic feature transcription of the first 2 phonemes of each word. These features describe the articulatory properties of every speech sound, based on the interactions of different physiological structures, and convey information about the broad phonetic category (whether the phone is voiced, unvoiced etc) and about place and manner for consonants (e.g., Nasal, Fricative, Coronal) and for vowels (e.g., Front, Close, Rounded). As discussed earlier, extensive multi-modal evidence supports a substantial role for such features in the early stages of speech analysis. Here we used standard binary articulatory feature transcriptions for each phoneme, as in Wingfield et al. (2017). A representational dissimilarity matrix (RDM) was calculated across the word stimuli, where the dissimilarity between phonological feature vectors (concatenated for the first 2 phonemes) was calculated for all stimuli pairs. A two-phoneme (as opposed to longer) model was chosen because of the focus here on the early stages of processing from word onset, prior to word recognition point. Most words have become identifiable by the time a third phoneme has been heard.

The *cohort entropy model* provides a measure of lexical uncertainty about the identity of the word being heard, based on the word candidates in the estimated cohort at each gate. This measure provides information about both the number of potential lexical forms and their likelihood, and was calculated with a modified version of Shannon’s entropy (H):

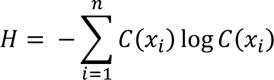

*where C(X_i_)* refers to the summed confidence score for a cohort competitor *i* across participants divided by the total sum of confidence scores for all the competitors, and *n* refers to the total number of competitors. High entropy indicates a large cohort with low confidence for any single candidate, and low entropy indicates few candidates with high confidence. The cohort entropy RDM was calculated as the absolute difference between the entropy values for each stimulus pair. This measure of lexical entropy may be related to, but is not identical to, the phoneme transition probability statistics (phonotactics) examined by Leonard et al (2015) in a previous ECOG study. For a fuller discussion of such relationships, see Gwilliams & Davis (2022).

The *cohort semantic model* captures the summed semantic information activated by the multiple word candidates across different gates. For each word candidate in the cohort, the semantics of that word was estimated using a corpus-based Distributional Memory database (Baroni & Lenci, 2010). The database represents the semantics of words as vectors over 5000 dimensions, where the entries of the semantic dimensions are derived from word co-occurrence data. The semantics of the whole cohort was calculated by averaging the semantic vectors of the candidates (each multiplied by the confidence scores), resulting in a 5000-dimension vector for each of the stimuli. The cohort semantics RDM was calculated as the cosine distance between the blended semantic vectors for each stimulus pair.

RDMs for cohort entropy and cohort semantics were calculated for cohorts established for three gates (50, 100 and 150 ms) covering the listeners’ processing of the relevant epoch after speech onset. The intended differentiation of the two types of model is reflected in the weak (though significant) correlation between them (r = +0.17, p <.001). In this respect these models are similar to the Entropy and Semantic Blend models constructed by Klimovich-Gray et al (2019) for two-word contexts, also weakly correlated at +0.17, and where each model generated very different patterns of model fit.

### Searchlight RSA

The relationship between the representational geometries of the cognitive RDMs and of the MEG signals was tested using spatiotemporal searchlight RSA (Su et al., 2012). Using the parameters estimated during source localisation to allow a mapping between each individual’s MEG sensor activity and the 5124 vertices on the canonical cortical surface, we can extract activity patterns for each trial within a spatial and temporal window (spm_eeg_inv_extract.m). At each vertex and time-point *t*, single-trial MEG signals were extracted for all vertices within a 10 mm radius for timepoints within a 64 ms time window centred at *t* (following Lyu et al., 2019). Multivariate noise normalisation (Guggenmos et al., 2018) was applied to these signals before RDMs were calculated as 1-Pearson’s correlation between all pairs of single trial responses. This procedure was repeated for all vertices and timepoints between −200 and 500 ms (at 4 ms increments).

To focus our analysis on regions known *a priori* to be involved in early phonological, lexical and semantic language processing, the searchlight analysis was restricted to a large bilateral fronto-temporal mask including bilateral inferior frontal (IFG), superior temporal (STG) middle temporal (MTG), supramarginal (SMG) and angular gyri (AG) (see Kocagoncu et al., 2017, Figure 2). The spatial definitions of these regions were taken from the Automated Anatomical Labelling (AAL) Atlas (Tzourio-Mazoyer et al., 2002), and were fused together as a contiguous mask with 1 mm isotropic spacing.

**Figure 2.**
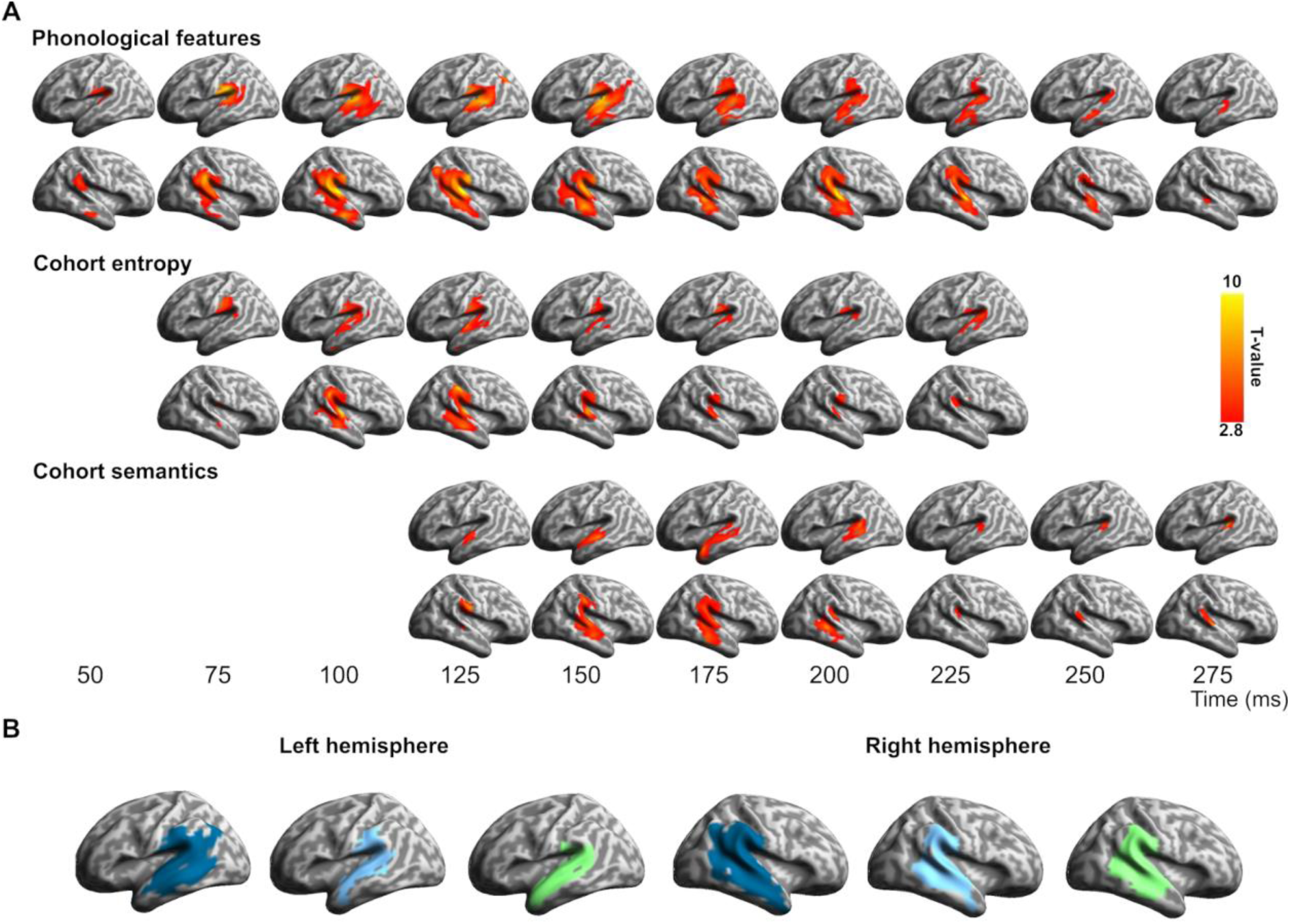
Spatiotemporal searchlight RSA results. A) Significant model fit for the phonological, cohort entropy and cohort semantics RDMs individually, plotted for datapoints spaced at 25 ms intervals (vertex-wise p < 0.01, cluster p < 0.05). B) Spatial extent of significant fits, collapsed over time, for phonology (blue), cohort entropy (cyan) and cohort semantics (green).

The MEG data RDMs for each point in space and time were compared against the dissimilarity values computed for the cognitive models using Spearman’s correlation, resulting in a Spearman correlation timeseries for each vertex and for every participant and cognitive model.

### Statistical analysis

Random effects analysis testing for positive RSA effects was conducted using a cluster-based permutation test (Maris & Oostenveld, 2007) performed on all cortical surface vertices within our language mask. To quantify the observed clusters, a one-sample t-test against zero (alpha 0.01) was performed at each vertex and for every timepoint. From this, clusters were defined by contiguous above-threshold effects over neighbouring timepoints and/or neighbouring vertices on the cortical mesh. The sizes of the observed clusters were calculated as the sum of the t-values for all time/spatial points within each cluster. To assess the statistical significance of the observed clusters, p-values were determined by creating a null distribution of cluster masses based on 1000 permutations of the MEG signals. For each permutation, the sign of the RSA correlations was randomly flipped for each participant, before conducting one-sample t-tests of the permuted data at each vertex and timepoint. The mass of the largest cluster (sum of above-threshold t-values) at any timepoint or location was retained for the null distribution. The cluster p-value for each of the observed clusters was defined as the proportion of the 1000 permutation cluster masses (plus the observed cluster-mass) that were greater than or equal to the observed cluster-mass. Anatomical labelling of the significant clusters was performed using the AAL Atlas for SPM (Tzourio-Mazoyer et al., 2002), where each vertex on the canonical mesh was assigned a label based on the corresponding MNI coordinates of the vertex and labels.

While the above analysis allows for inferences about the presence of effects, it does not allow for inferences about the relative onsets of effects for different model RDMs (Maris & Oostenveld, 2007; Sassenhagen & Draschkow, 2019). To test for differences in the onsets of effects, a jack-knife resampling procedure was used (Miller et al., 2009). Using the RSA time series at individual vertices, onsets were calculated for each of the jack-knife resampled datasets as the first time-point (post 0 ms) with a significant p-value using a one-sided t-test against zero, creating a distribution of onset times for each model RDM. A sign-rank test was then used to compare between the onset distributions of different model RDMs to determine if onsets were reliably earlier for one model RDM compared to another.

## Results

To examine the spatiotemporal evolution in real time of the early analyses of spoken words heard in isolation, we obtained measures based on the initial two phonemes of the word (phonological features), the early lexical candidate set (cohort entropy), and the blended semantic representations of these lexical candidates (cohort semantics). Cohort entropy and cohort semantics models were calculated for each of three gates at 50, 100 and 150 ms. Representational dissimilarity matrices (RDMs) based on these measures were tested against source reconstructed MEG data RDMs using spatiotemporal searchlight RSA. An overview of the RSA searchlight effects (Figures 2 and 3), describing the overall pattern of the results, is followed by detailed analyses of the spatial and temporal profile of the neural model fit for each model tested (Figures 4, 5 and 6, and Table 1).

**Figure 3.**
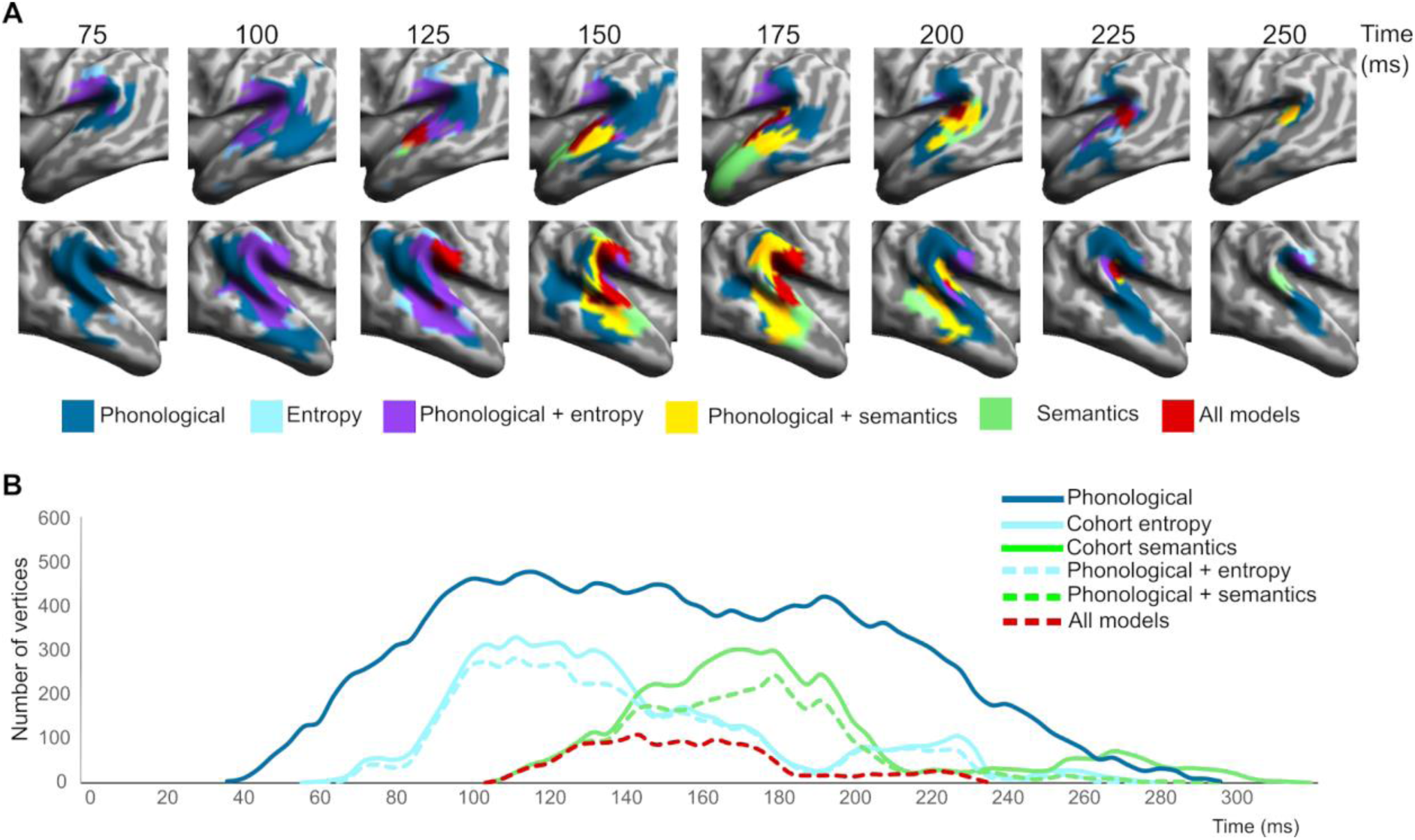
Spatial and temporal overlap of significant vertex-wise RSA model fit across models. A) Spatial overlap of different model fits over the test epoch in core temporo-parietal regions, shown for data points spaced at 25 ms intervals. B) Temporal distribution of vertices (sampled at 4 ms intervals) showing different types of model fit, including (i) overall model fit (unbroken lines) for phonology (dark blue), entropy (cyan), and semantics (green), (ii) vertices showing significant model fit to both entropy and phonetics (broken cyan line), (iii) vertices showing model fit to both semantics and phonetics (broken green line), and (iv) vertices showing fit to all three models at the same time (broken red line).

**Figure 4.**
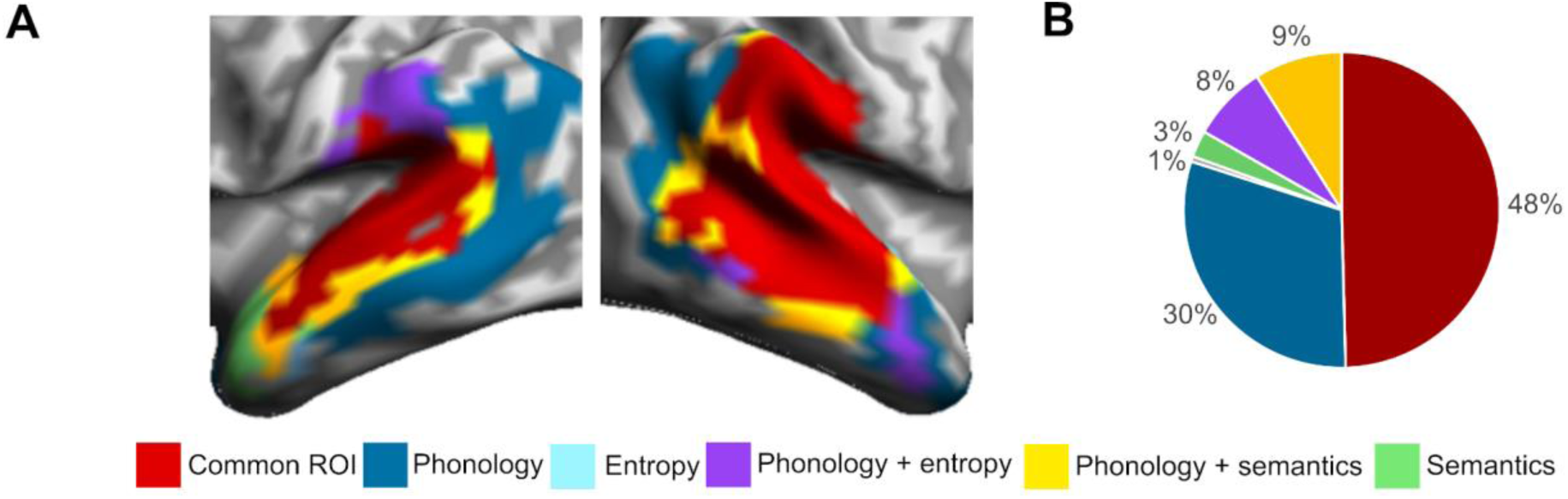
Spatial overlap in commonly activated vertices across model fits. A) Overlap in spatial extent of significant fits for each model RDM, collapsed over time. B) Percentage of vertices that show common effects for all three models, for two models or for a single model.

**Figure 5.**
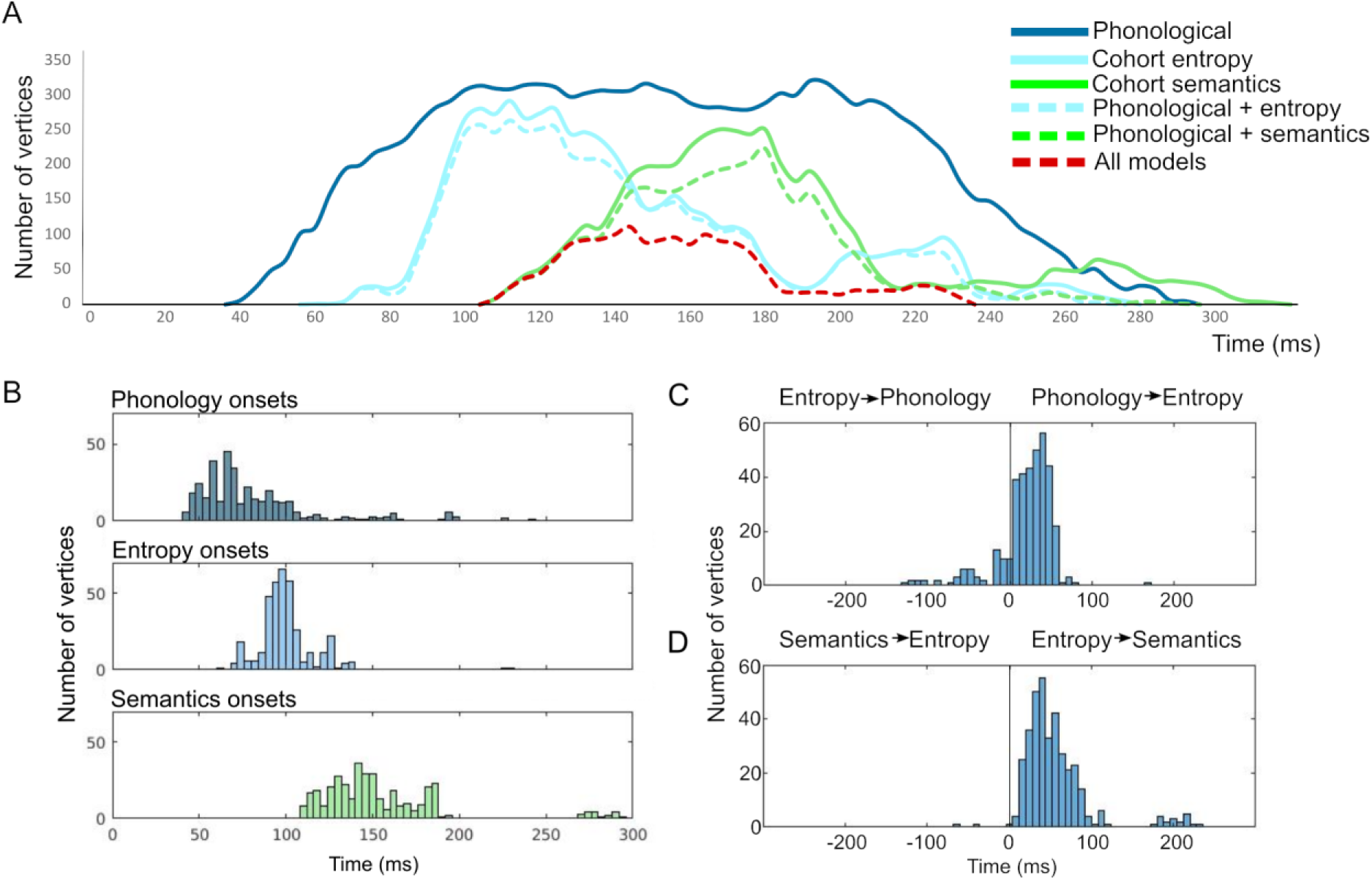
The temporal progression of effects within the common ROI. A) Temporal distribution of vertices within the common ROI showing different types of model fit (sampled at 4 ms intervals). B) Histograms of the onsets of model significance at each vertex within the common ROI. C) Distributions of temporal delays between phonology and entropy model fits for each vertex, and D) Distributions of temporal delays between entropy and semantic model fits for each vertex.

**Figure 6.**
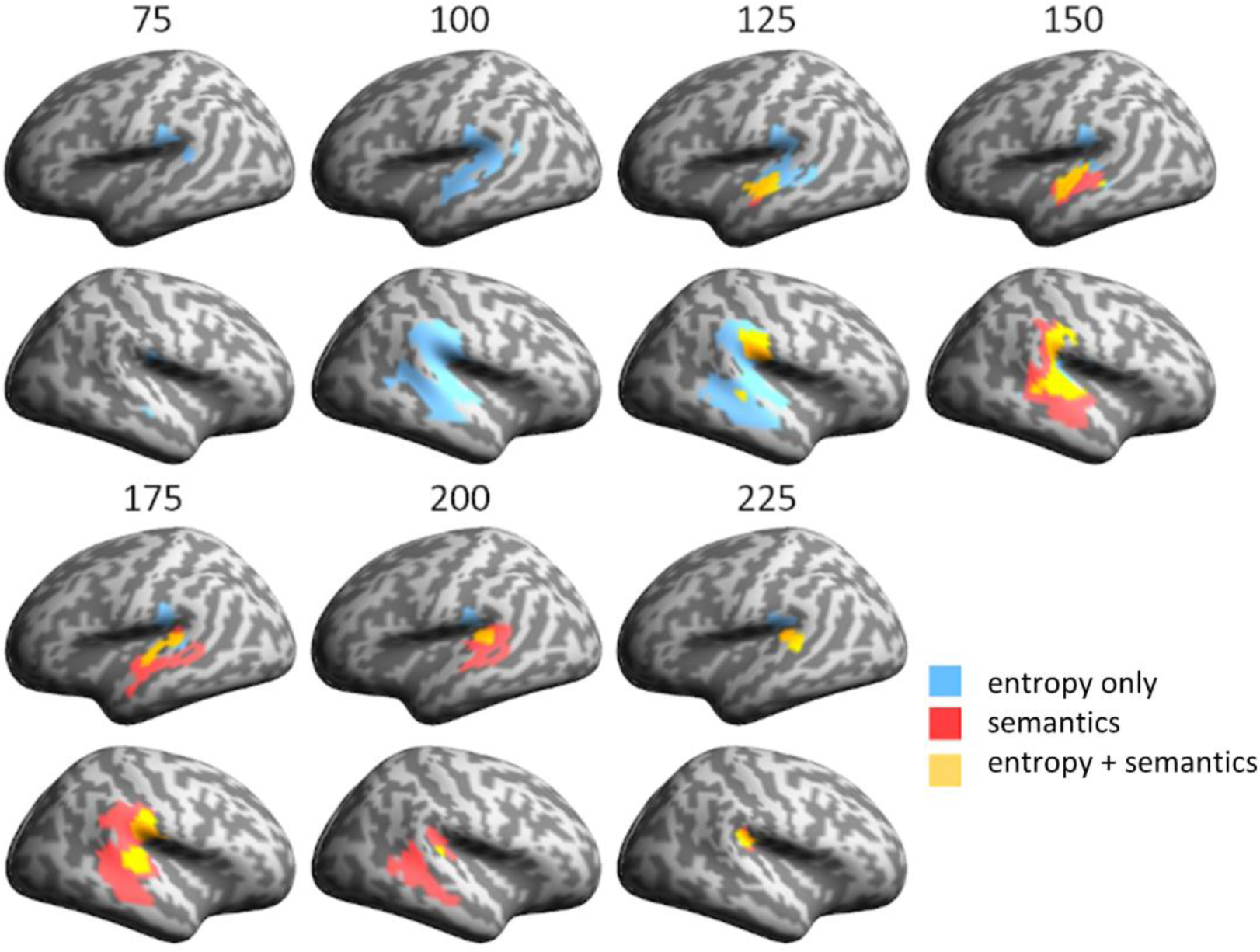
Spatiotemporal relationship of entropy and semantic model fits within the common ROI: Vertices that are initially significant to entropy only (blue) become either simultaneously significant to entropy and semantics (yellow) or become significant to semantics only (red).

**Table 1.** Distribution of vertices across anatomical regions (AAL atlas). RSA model fit (in number of vertices and as a percentage of ROI total vertices) is shown for the common ROI and for each model separately. Total size (in vertices) of each AAL region is given in the left-hand column. AAL regions with model fits involving less than 10 vertices or less than 10% of ROI are not shown.

| Regions | Size | Phonology |  | Entropy |  | Semantics |  | Common |  |
| --- | --- | --- | --- | --- | --- | --- | --- | --- | --- |
|  |  | N. vert | % ROI | N. vert | % ROI | N. vert | % ROI | N. vert | % ROI |
| L STG | 81 | 62 | 77% | 68 | 84% | 70 | 86% | 56 | 69% |
| R STG | 114 | 102 | 89% | 97 | 85% | 106 | 93% | 92 | 81% |
| L MTG | 166 | 136 | 82% | 53 | 32% | 82 | 49% | 48 | 29% |
| R MTG | 135 | 111 | 82% | 68 | 50% | 74 | 55% | 52 | 39% |
| L SMG | 52 | 46 | 88% | 33 | 63% | 10 | 19% | 9 | 17% |
| R SMG | 91 | 83 | 91% | 79 | 87% | 77 | 85% | 76 | 84% |
| R inferior parietal lobe | 52 | 8 | 15% | 2 | 4% | 4 | 8% | 2 | 4% |
| L angular gyrus | 50 | 34 | 68% | 0 | 0% | 0 | 0% | 0 | 0% |
| R angular gyrus | 72 | 42 | 58% | 0 | 0% | 1 | 1% | 0 | 0% |
| R Heschl gyrus | 11 | 7 | 64% | 7 | 64% | 7 | 64% | 7 | 64% |
| R rolandic operculum | 61 | 11 | 18% | 11 | 18% | 11 | 18% | 11 | 18% |
| R inferior temporal lobe | 92 | 12 | 13% | 2 | 2% | 6 | 7% | 1 | 1% |
| L superior temporal pole | 20 | 0 | 0% | 0 | 0% | 7 | 35% | 0 | 0% |
| R superior temporal pole | 24 | 4 | 17% | 1 | 4% | 1 | 4% | 0 | 0% |
| L middle temporal pole | 14 | 0 | 0% | 0 | 0% | 2 | 14% | 0 | 0% |
| R middle temporal pole | 18 | 15 | 83% | 0 | 0% | 0 | 0% | 0 | 0% |
| Total number of vertices |  | 710 |  | 446 |  | 471 |  | 364 |  |

### Early model fit and spatiotemporal overlap

The salient overall result is the rapid sequential emergence of model fit for all three models underpinned by a strong spatial and temporal overlap in the central neural expression of these effects (Figure 2). The Phonological Feature model (Figure 2A) showed the earliest and most extensive fit to the brain data, starting within approximately 40 ms of word onset in posterior superior temporal gyrus (STG) and rapidly engaging an extensive bilateral network including superior and middle temporal cortex and extending into anterior temporal and inferior parietal regions, including angular gyrus and supramarginal gyrus (Left hemisphere: cluster p < 0.001, time window = 44-292 ms; Right hemisphere: cluster p < 0.001, time window = 40-292 ms). This spatiotemporal pattern is consistent with neurobiological models of speech interpretation which predict a largely feedforward flow through temporal and parietal cortex, and is compatible with timings from earlier studies of phonological processing (e.g. Brodbeck, Hong, et al., 2018; Brodbeck, Presacco, et al., 2018; Broderick et al., 2019; Di Liberto et al., 2015; Mesgarani et al., 2014; Travis et al., 2013).

Looking beyond phonological properties, however, the results challenge a spatially segregated account where different neural regions are tuned to different aspects of early speech analysis. Spatiotemporally closely related to the phonological results, the RSA analyses picked out significant but more transient effects for the two cohort-based RDMs (Figure 2A), both of which generated significant model fit only for the RDMs using the gate 150 ms candidates (Cohort Entropy: Left hemisphere: cluster p = 0.004, time window = 60-240 ms; Right hemisphere: cluster p < 0.001, time window = 68-276 ms; Cohort Semantics: Left hemisphere: cluster p = 0.006, time window = 108-306 ms; Right hemisphere: cluster p = 0.003, time window = 108-296 ms). The striking degree of overlap between the three models in the spatial distribution of model fit is illustrated in Figure 2B. The absence of significant effects for the 50 and 100 ms gates is likely to reflect the instability of lexical choices at these earlier time-points.

The spatial and temporal relationship between the three RDMs is examined in more detail in Figure 3, with Figure 3A showing the evolving pattern of spatial overlap between models over the first 300 ms, while Figure 3B focuses on the temporal unfolding of this process, showing the relative timing of model fit for each RDM. Early bilateral model fit for the phonological model is followed within 25-30 ms by cohort entropy model fit around 60-70 ms post-onset, engaging largely the same vertices in the same regions of superior and middle temporal cortex (Figure 2B, Figure 3A). Entropy model fit continues to track the phonological model as it expands anteriorly and posteriorly, engaging bilateral supramarginal (but not angular) gyri. The spatial extent of entropy model fit reaches its peak around 100 ms post word onset and begins to drop away after around 140 ms (Figure 3B, solid cyan line). Saliently, the great majority (94%) of entropy sensitive vertices also show significant phonological model fit (Figure 3B, broken cyan line). The timing of these entropy effects corresponds closely to the early lexical entropy and related phoneme surprisal effects reported in previous studies (for a summary see Gwilliams & Davis, 2022, Figure 5.4), even though almost all these studies (as noted previously) involved continuous speech rather than isolated words.

The cohort semantics model shows a very similar overall pattern (Figure 2B, 3A), overlapping with both phonological and entropy models in superior and middle temporal cortex, most strongly in the right hemisphere, but displaced by 40-50 ms relative to cohort entropy. The spatial extent of model fit for cohort semantics peaks near 180 ms followed by a steep drop off after around 200 ms (Figure 3B, solid green line). Similar to cohort entropy, the great majority of vertices (91%) sensitive to cohort semantics show model fit to the phonological model (Figure 3B, broken green line). In terms of the processing relationship between the two cohort variables, 86% of vertices sensitive to entropy also show an effect for semantics, while 82% of vertices sensitive to semantics show an effect for entropy. Note that all vertices shown as sensitive to both entropy and semantics at the same time were also sensitive to phonology (Figure 3B, broken red line).

These analyses show that the early stages of access to lexical form and meaning from phonological cues in the speech input engage a common network of shared model-sensitive vertices in bilateral anterior and posterior superior and middle temporal cortex, and right supramarginal gyrus. Critically, this network does not seem to be segregated into different subregions supporting different aspects of the early access process. We now turn to a more specific spatial and temporal definition of the properties of this candidate core network.

### Defining a common core

We define the common network as including all (and only) those vertices that showed significant model fit to all three model RDMs, at any point in time over the test epoch (see Figure 4 and Table 1). Because this definition of the common network collapses over time (similarly to Figure 2B), it will include vertices where the model fits to different models may occur sequentially as well as simultaneously. In terms of the spatial distribution of this network, Figure 4A plots the spatial overlap in model fit between the three models, separating the common ROI (red) from surrounding regions where model fit was either seen for just two of the models (phonology and entropy, phonology and semantics), or for one of the models individually. Figure 4B summarises the distribution of vertices across these alternatives. Table 1 gives the general location of significant vertices, for each model RDM and for the common network, in relation to the principal anatomical regions involved.

The common ROI captures 48% of all the vertices in the bilateral language mask that were significant for any model (Figure 4B) and is largely (90%) clustered in 3 core bilateral temporo-parietal regions (STG, MTG, and SMG), most strongly in STG and right SMG (Table 1). This network extends partially into bilateral MTG but only weakly into left SMG, with a strong bias overall to the right hemisphere (68% RH vs 31% LH). Model fit for the Entropy and Semantics RDMs closely follows the pattern for the common ROI with each showing 89% of significant vertices falling within the core temporo-parietal regions and with the highest density of model fit in bilateral STG and right SMG (Table 1). Both models show the same imbalance in favour of the right hemisphere (61:36% for Entropy and 61:35% for Semantics).

Model fit for the Phonological model stands in a different relation to the common ROI, relative to the Entropy and Semantic models, by virtue of its greater spatiotemporal extent, covering 710 vertices. Half of these vertices (51%) fall within the common ROI, showing extensive model fit for the core bilateral temporo-parietal regions, though with much increased model fit in left SMG and bilateral MTG (Table 1; Figure 4A). Extending well beyond the common core, the Phonological model also shows substantial shared model fit to each of the other models separately (Figure 4B) - with semantics in middle and posterior temporal regions bilaterally, and with entropy in left SMG and right posterior temporal regions. A further 30% of vertices show phonological model fit uniquely, most notably in the angular gyrus, extending into posterior STG bilaterally and left MTG. The absence of any lexical model fit for these latter areas suggests a quite different role in the early speech interpretation process – possibly reflecting a dorsal prosodic route (Hickok et al., 2022; Hickok & Poeppel, 2015). Finally, the temporal duration of phonological model fit, together with its extensive spatial coverage, means that almost all the vertices involved share a uniform and stable phonologically defined representational geometry throughout this early speech analysis period.

### The evolving representational geometry of the common core

Given this combined anatomical and functional specification of a potential common processing core, we can then determine the nature of the temporal progression whereby the vertices in this network show model-fit to the three different model RDMs. Figure 5A shows the relevant temporal distribution of vertices, restricted to those within the common core. Examining the temporal dynamics of model fit for each vertex within the common network (Figure 5B), we find that 64% of the vertices have a significantly earlier onset for phonology than entropy (all p < 0.05; mean delay = 20 ms, mode = 44 ms; Figure 5C), and that 75% of vertices show a significantly earlier onset to entropy compared to semantics (all p < 0.05; mean delay of 40 ms, modal = 44 ms; Figure 5D). But to understand how the representational geometry of the common core evolves over time, we also need to know how far model fit for different models shows a sequential relationship (e.g. model A is significant, then model B is significant for the same vertices but with no temporal overlap), or shows at least partial overlap in time, whereby the vertices are simultaneously significant for both (e.g. model A is significant then model A+B are both significant).

A preliminary answer emerges from the overall temporal distributions plotted in Figure 5A. These confirm that model fit to the Phonological RDM is dominant throughout this early epoch (∼60-250 ms), reflecting the stability during this period of a representational geometry capturing the speech perceptual input (seen from an articulatory perspective). Superimposed on this backdrop are the temporally staggered bursts of model fit for the Entropy model (peaking between ∼80-140 ms) and for the Semantic model (∼150-200 ms), each encoding a representational geometry that is only transiently congruent with the listeners’ incremental brain states as the word was heard. This means that all vertices showing Entropy or Semantic model fit can in principle be simultaneously sensitive to Phonology, whereas for many vertices the relationship between Entropy and Semantics model fit will necessarily be sequential, since Entropy model fit is already peaking at 100 ms before Semantic model fit has even begun (see Figure 5A, B).

A more detailed vertex-wise analysis, also restricted to members of the common ROI, confirms this general picture of the evolving pattern of representational onsets and offsets. As noted earlier, the great majority of these vertices show a simultaneous relationship between phonology and entropy (93%), and between phonology and semantics (82%), but this drops to 58% for entropy and semantics. Here the number of vertices showing a sequential relationship for entropy and semantics markedly increases to 42%.

This increase in the proportion of vertices showing sequential model fits seems to follow from the temporal delays in model fit between the Entropy and Semantic RDMs. As Figure 5D documents, almost all Common ROI vertices start to show Entropy model fit before they show Semantics model fit (at a mean delay of 40 ms), presumably reflecting the uniform evolution of brain representational geometry as the speech input accumulates. This generates a gradient shift in model fit, propagating across the common ROI, from the entropy RDM to the semantics RDM. But because of the steep drop off in Entropy model fit from around 140 ms, only the earlier vertices showing Semantic model fit can do so while still showing Entropy fit. These are the vertices showing ‘all model’ fit in Figure 5A and are a subset of the vertices showing early Semantics model fit in Figure 2A. Direct evidence for this is the close overlap in Figure 5A between the red ‘all models’ plot and the first 35-40 ms of semantic model fit (unbroken and broken green lines), starting at around 100 ms.

In Figure 6, we examine in more detail the spatial overlap between entropy and semantic model fits across vertices within the common ROI (excluding phonological model fit from the visualisation). This shows how entropy model fit (peaking between 100-140 ms) is gradually replaced by semantic model fit (peaking between 150-200 ms) as it propagates across the common ROI. The earliest semantic model fit emerges in bilateral posterior STG and right SMG. At these early time points, fit to the two model RDMs largely co-occurs (coded in yellow in Figure 6) and is maintained as such until around 175 ms, by which time entropy model fit is minimal (Figure 2A, 5A). Sequential model fit (coded in red) becomes more dominant from around 150 ms, defined as cases where vertices initially sensitive to entropy later become significant only for semantics (while maintaining model fit to phonology throughout). These later fitting vertices are primarily located in bilateral MTG and in more posterior right perisylvian cortex.

In summary, our results support a spatially homogenous process of early speech analysis, distributed across bilateral temporoparietal brain regions, where the process of incremental interpretation generates spatiotemporally evolving representations whose primary geometry is captured by the dissimilarity structure of the phonological input, with this representational structure being maintained and rendered more lexicosemantically specific as accumulating acoustic-phonetic constraints propagate across the common ROI and beyond.

## Discussion

This research addresses two foundational questions about the earliest stages of speech comprehension. These concern the functional and neurobiological architecture that underpins these processes and the computational vocabulary in terms of which they are conducted. In doing so, we contrast the widely held representational hierarchy model with a fully distributed network approach, viewed against the backdrop of extensive ECOG research pointing to a potentially intermediate view, where early speech analysis is subserved by a distributed mosaic of specialised processors, interpreting different acoustic-phonetic cues in the speech signal.

To address these issues, we used ssRSA of source-localised MEG signals to provide a neurocomputationally specific spatiotemporal map of the emergence of three critical dimensions underlying the early stages of speech interpretation - the phonological integration of acoustic-phonetic information in the speech signal, the implications of this information for potential lexical identities intended by the speaker, and the access of semantic information related to these early cohort members. These analyses provide a well-specified regional map of the distribution of these three processing dimensions, tested within a broad fronto-temporo-parietal language mask. Because of the nature of the MEG signal, the properties of the source localisation process, and the size of the searchlight window, this is likely to be a low-dimensional mesoscale view of the population dynamics of the neural activity underpinning speech interpretation. While the mechanistic basis for such a causal cortical level of analysis is not well understood (c.f. Pinotsis & Miller, 2022), it is widely accepted that the representational geometry of mesoscale neural populations is essential to the explanation of complex neurocognitive functions (Barack & Krakauer, 2021; Chang, 2015; Chung & Abbott, 2021; Kriegeskorte & Kievit, 2013). In so far as representational geometry is intrinsically a mesoscale property of the cortical fields generated during speech interpretation, then the MEG/ssRSA method is likely to be well adapted to tracking the spatiotemporal dynamics of such geometries.

This mesoscale view of likely MEG/RSA model fit brings with it potential caveats about the spatial resolution of these techniques relative to the questions we are asking. This is unlikely to be a problem, however, in evaluating the primary theoretical contrast between a distributed DISCO-type model and a stratificational hierarchy locating phonemic analyses in superior temporal cortex and lexically based analyses in middle temporal cortex. There is ample evidence that ssRSA model fit in MEG source space can be sufficiently neurocomputationally specific and spatially well-defined to tell a convincing story about the patterning of different neurocognitive process as they evolve over time in these core temporal language areas.

We see this, for example, in the Kocagoncu et al (2017) RSA analyses for the same dataset, which showed well circumscribed and differentiated patches of model fit for the different lexical and semantic models tested, while Lyu et al (2019) use MEG/ssRSA to reveal in unprecedented neurocognitive detail the neural dynamics of lexical semantic interpretation in sentential contexts. In earlier papers focusing on acoustic and phonological processes in medial and lateral STG, we use MEG/ssRSA techniques to pick out spatiotemporally well-defined patches of model fit corresponding to different subsets of auditory input analyses (Su et al., 2014; Wingfield et al., 2017, 2022). For the analyses reported here, there are several examples of spatially differentiated model fit – for example left and right angular gyrus and right middle temporal pole all show strong fit to the phonology model but no fit to the entropy or semantics models (Figure 3 and Table 1).

At the same time, however, we cannot rule out *a priori* a processing scenario where a sufficiently fine-grained mosaic of independent processors could jointly contribute to the representational geometries measured at each searchlight testing point. But this is not a scenario we have seen proposed anywhere in the stratificational literature. Nor is it obvious how a stratified phoneme-based model could be mapped onto a mosaic of localised independent processors linked to the phonological, lexical, and semantic properties of speech analysis, with the constraint that these processes are evenly distributed across the common ROI (as identified in the current paper), and where the computational autonomy of each type of process was still preserved. In the absence of such a model, we will focus here on the conventional hierarchical model of spatially separated computational domains allocated to different sulci and gyri across superior and middle temporal cortices.

Bearing these concerns in mind, the resulting set of maps reveal a stable bihemispheric core of common processing activity, centered around bilateral posterior temporal and parietal cortex, where the same set of vertices exhibit, simultaneously and in sequence, sensitivity both to the phonological structure of the speech signal and to the lexicosemantic implications of this structure, with no apparent spatial segregation of these different types of neural content. The absence of any contextual constraint for this set of isolated words means that the observed co-location of these early model fits cannot be attributed to external tuning of the processes involved. Instead, the results reflect the intrinsic bottom-up processing architecture of the system.

### The neurocomputational environment for early speech analysis

What is the global processing environment revealed by these patterns of model fit? The overarching representational geometry that dominates early speech processing seems well captured by the phonological feature RDM (Figure 3B). This RDM is constructed from the concatenated articulatory feature decomposition of the first two phonemes of the word being heard and remains constant throughout the test epoch. As noted earlier, we view this model as a proxy for the articulatory gestures that generate the spectrotemporal modulation of the speech waveform over time. The presence of substantial model fit to this dissimilarity matrix for over 200 ms from word onset means that the primary structure of the neural geometry being tested, viewed from this articulatory perspective, will also remain stable over this early epoch. This stability reflects, in part, the structuring of the perceptual analysis space by the earliest arriving acoustic-phonetic information about the incoming word, consistent with the cohort-based approach to speech interpretation (Marslen-Wilson, 1987). It may also reflect the longer-term maintenance across the common core (and beyond) of the neural substrate for the listener’s perceptual experience of the speech they are hearing.

Within these constraints, however, this neural geometry will necessarily evolve as the epoch progresses, responding to the accumulating acoustic-phonetic information carried by the speech stream. For the word *bat*, for example, the space of lexical possibilities is defined within 50 ms of word onset by an initial articulatory gesture that is bilabial, voiced, and obstruent. This primary structure is gradually modulated by cues to the properties of the following vowel – here an [ae] - which itself begins to provide cues to the following articulatory target – in this case coronal, unvoiced and obstruent – with the intended word emerging as the perceptual outcome. This evolution within the early epoch towards greater lexical specificity does not, however, seem to disrupt the basic representational geometry of the speech perceptual space, as indexed by the continued strength of phonological model fit over this period.

The properties of this evolving neural geometry are illuminated by the timing and the computational content of the model fit shown by the two lexically related model RDMs (entropy and semantics). Both have the properties of *transienc*e and of *sharing model fit* with each other and with the phonology RDM for all vertices within the common core. This three-way shared model fit strongly suggests that we are not dealing here with three separate representational model spaces, but rather a single integrative representational space. The transience of entropy and semantic model fit means, therefore, that these are snapshots, at different points in time, of the properties of this space as it evolves towards a unique perceptual solution as the word is recognized (Kocagoncu et al., 2017).

Within 20-30 ms of the first emergence of phonological model fit, we start to see model fit for the lexical entropy RDM, where this encodes a dissimilarity structure which captures the distribution of lexical uncertainty across the stimulus set. The fact that this model tracks so closely in time and space to the patterning of early phonological model fit suggests that the lexical implications of the articulatory events encoded by the phonological RDM are integral to the speech analysis process, even at these very early stages. There were no contextual constraints that could provide top-down cues to potential lexical candidates, and it is implausible that phoneme sequence constraints (e.g. Gwilliams et al., 2018, Figure 8) could interact with potential lexical candidates in MTG early enough to generate lexical entropy effects at the timing we observe.

Entropy model fit is nonetheless much more transient than phonological model fit, peaking at around 100 ms post onset and dropping away sharply 40-50 ms later. This means that there is only a short period within which the dissimilarity matrix generated by variations in lexical entropy comes into register with the primary geometry of the brain’s early interpretation of the speech signal, as captured by the phonology RDM. Instead, the cohort semantics RDM takes over, most strongly from around 140 ms to 200 ms, indicating a period during which the dissimilarity matrix reflecting the semantic clustering of word candidates comes into register with the same underlying neural geometry. The transience and the co-location of these model fits makes it hard to see how they can have their own ‘representations’ – that is, as reflecting separate bursts of neural computation specific to analyses of lexical form and lexical meaning. Instead, they reflect the evolving representational geometry of the common core - captured by the articulatory feature RDM and enriched by greater lexicosemantic specificity as further constraints accumulate. Critically, this lexicosemantic ‘enrichment’ must be generated internal to the analysis process, since there is no external context to provide cues to particular forms and meanings.

These considerations suggest that the perceptual analysis being computed during early speech processing is one that synthesizes together constraints from multiple population-coded aspects of the speech interpretation process. A primary function of this process is to generate the listener’s phenomenological experience of what is being said. This is necessarily a speech percept, but it is not simply a record of the listener’s sensory experience, it is a linguistically interpreted speech percept. The groundwork for this is laid during these early, first pass integrative analysis processes.

### Implications for the architecture of early speech interpretation

The picture of early speech analysis that emerges from the pattern of ssRSA model fits is markedly problematic for the conception of this process as a hierarchical sequence of representationally distinct stages of analysis across different sulci and gyri. Most obviously, there is a major macroscale discrepancy between the global neural architecture required by a sequential hierarchy and the architecture we actually observe.

To manage the translation from one level of representation to another in a sequential hierarchy, the proposed levels of analysis are standardly realized as spatially and temporally segregated levels of neurobiological process. In line with this, current hierarchical models postulate an anatomically and computationally distinct layering of primarily feedforward processes, radiating from spectrotemporal analyses in Heschl’s gyrus to phonetic feature extraction in STG, leading to phoneme identification and thence to potential lexical candidates in MTG (e.g. DeWitt & Rauschecker, 2012; Gwilliams et al., 2018; Hickok & Poeppel, 2007). The ssRSA picture looks nothing like this. Instead, we see a substantial common core of vertices, covering many of the same areas bilaterally, that exhibit sensitivity to aspects of early speech interpretation that cut across all these domains, but with no sign of neuroanatomical or neurocomputational segregation. The spatial co-location of these disparate neurocomputational functions is directly inconsistent with the defining properties of a layered hierarchical approach.

A particular difficulty, as noted earlier, is the prevalence of three-way model fit (predominantly simultaneous) to each common core vertex by three apparently orthogonal models. In a conventional hierarchical representational system, it is not possible for the same vertex (let alone a whole contiguous region of vertices) to simultaneously encode theoretically distinct processing operations. To do so violates the foundational requirement for both anatomical and computational segregation of the different representational domains distinguished by the underpinning hierarchical theory of the domain in question. Such an approach cannot accommodate a single integrative representational space within which the process of speech analysis might be conducted. It is this representational space that the combination of MEG source-localised data and ssRSA analyses seem able to tap into.

A final problem is the apparent incapacity of a sequential hierarchy to explain the critical role of speech information in constructing the listener’s speech perceptual experience. The processing goal of a conventional layered linguistic hierarchy is to achieve a transformation in what is being represented across successive levels. This process of sequential translation means that phonological analyses are discarded once they have fulfilled their role in mediating access to lexical form and meaning. The ssRSA analyses reported here show that the computational background for early speech analysis – spreading across substantial areas of bilateral temporoparietal cortex - is characterised throughout by the articulatory feature RDM, preserving the phonological essence of the message being communicated.

A fully distributed network approach, in contrast, provides a natural fit to these overlapping spatiotemporal patterns of neural computation. Such an account does not need to impose a top-down conception of the specific sequential structure and content of the analysis process, and then try to fit the data to the neurocomputational categories entailed by this structure. Instead, it assumes that the distributed architecture of the human brain finds its own, *sui generis* solution to the problem of mapping from speech input onto an integrative perceptual interpretation, and asks what the properties of this solution might be, in terms of its structure and neurocomputational content.

This was the starting point for the Distributed Cohort (DisCo) concept (Gaskell & Marslen-Wilson, 1999, 1997), which rejected a sequential ordering of discrete processing levels and argued instead for a view of speech perception as a direct mapping from low-level featural information onto a distributed representation of lexical knowledge and form (as proposed by Marslen-Wilson & Warren, 1994). In the original DisCo implementation, this mapping was handled within a single fully distributed representational space, based on a simple (shallow) recurrent network architecture (Elman, 1990) such that the identification of a word involved the simultaneous integration of both phonological and lexicosemantic constraints. More generally, we assume, a distributed recurrent architecture of this sort does not need to be specialised for particular qualitative types of analysis at these early processing stages, translating between successive intermediate representational steps. It should be seen instead as a complex and effective distributed mechanism for learning how to compute an optimised pathway through the neural maze to a perceptually interpreted outcome. In doing so it will necessarily encode the statistical regularities underpinning this mapping but without needing to segment these regularities into distinct computational types (though see Stephenson et al, 2020).

These hypothesized properties of the DisCo type of architecture are consistent with just those aspects of the current results that are most problematic for a layered hierarchy – namely, the co-location of model fit at the same vertices, realised as the sequential but transient appearance of lexical entropy and lexical semantic model fit in the overarching context of continuing articulatory feature model fit.

First, if early speech analysis is conducted in a distributed neurocomputational space, then the regions of vertices constituting this space will continue to be active as long as an analytic path is being computed through this space in response to a continuing sensory input. Second, as this dynamic solution evolves towards greater perceptual and lexicosemantic specificity, this will be reflected in the representational geometry of the neural space being evaluated. From very early in the analysis process, the learned weighting of paths through the network will reflect their relevance to the system’s integrated phonological and interpretative output goals. These changes, spreading across the common computational space as new constraints emerge, are mediated by the same neural substrate throughout, and are picked up, successively, by the lexical entropy and the lexical semantic RDMs. The transience and the relative timing of these lexically related model fits reflects this dynamic process of perceptual and interpretative refinement.

Thirdly, because the overarching neural geometry nonetheless remains defined in phonological terms, what is being computed preserves the basis for the listener’s speech perceptual experience. The observed continued activity across vertices, and their continuing adherence to a common representational geometry, suggests that what we are tapping into here is the neural substrate for the emergence and the stability of the listener’s dynamic speech percept.

More evidence for this comes from the breadth of model fit to the articulatory phonology RDM. On current views of early speech analysis (e.g. Bhaya-Grossman & Chang, 2022), middle and posterior STG are the main site for neural processes relating acoustic-phonetic cues to phonological structure. Here we see a match between the phonological structure RDM and patterns of neural activity in temporal and parietal regions far removed from STG (see Table 1 and Figure 4A), with around 50% of vertices sensitive to the phonology RDM falling outside the common core (and only 27% of them located in bilateral STG). Given our emphasis on the role of the speech processing network in generating the listener’s perceptual representation of speech, this wide distribution of phonological model fit reflects the dynamic interweaving of the perceptually salient phonological interpretation of the unfolding word with early emerging information about its interpretative implications. Yi et al (2019) have argued that STG plays a central role in combining multiple sources of speech related constraint into a unified perceptual representation. Our results suggest that this unification of multimodal inputs will also include core aspects of meaning, and will extend well beyond the STG.

### Relationship to ECOG research

Finally, we need to relate the proposals emerging here to the benchmark results generated by research using ECOG methods, revealing the neural organisation of early speech analysis in unprecedented electrophysiological detail. In the current context, this research stands in an intermediate position, between a linguistically stratified analysis process and a representationally neutral distributed process. Earlier ECOG studies (e.g. Canolty et al., 2007; Chang et al., 2010; Flinker et al., 2011) did adopt the dominant stratified approach, based on a linguistic array of intermediate processing types that define the progression from sound to meaning. More recent studies, focused primarily on left hemisphere STG and medial auditory cortex, have rejected a conventional sequential hierarchy, but are still committed to the notion of intermediate processors labelling and categorising a range of different cues which must then be integrated at a higher intermediate level.

Accordingly, confronted with the electrophysiological realities of STG speech analysis, current ECOG research has moved to a less hierarchical but still representationally segmented concept of STG, viewing it as a mosaic of multiple specialised ‘processing units’ (Bhaya-Grossman & Chang, 2022; Hamilton et al., 2021; Yi et al., 2019). These hypothetical units, identified probabilistically from the stimulus response profiles of individual ECOG electrodes, are argued to provide information in parallel about several intermediate stimulus properties relevant to speech interpretation. These include phonetic features (Fox et al., 2020; Mesgarani et al., 2014; Oganian et al., 2022), the temporal structure of the waveform (Hamilton et al., 2018; Oganian & Chang, 2019), and variations in relative pitch (Tang et al., 2017). While there is some clustering of speech-onset detectors in posterior and middle STG, electrode sites sensitive to phonetic features and to syllable onsets (‘peakRate’) are distributed across the entire STG (Hamilton et al., 2021).

At the same time, as Bhaya-Grossman & Chang (2022, p.84) point out, no cortical area was found that selectively encoded single phonemes or syllables at local sites. The Mesgarani et al. (2014) study, covering all the phonemes in English, showed that the different commonalities across speech-sensitive electrode responses were solely in terms of the articulatory features they shared. No selectivity for single phonemes was found. Similarly, the Oganian et al. (2022) study focusing on the STG substrate for vowel discrimination (in Spanish) found no evidence for focal representation of single vowels. Decoding of single vowels emerged robustly only at the population level, when information from all the differently tuned electrode sites was pooled together.

These properties of STG speech analysis, seen as a distributed network of probabilistically defined partially overlapping processes for extracting multiple constraints on speech interpretation, are well suited to a distributed recurrent network approach – either as generating the inputs to such a network or as a reflection of its dynamic operations in action. At the same time these properties are transparently inconsistent with a classic sequential hierarchy. Reflecting this, both Yi et al. (2019) and Bhaya-Grossman & Chang (2022) propose recurrent computational models of STG speech analysis – though see Keshishian et al. (2023) for a more hierarchical ECOG-based approach.

The main point of divergence between the DisCo approach and current ECOG theorising is not, therefore, the ground-breaking data that ECOG methods provide about STG function but rather the representational assumptions each approach makes in interpreting these data. ECOG models depend on specialised intermediate neural processes that label and categorise a range of different cues which must then be integrated at a higher level, leading to an architecturally and logistically complex set of further processes, especially where the role of phonemes is concerned. One example is the additional machinery discussed by Yi et al (2019, pp 1103-1105) for keeping track of the different phonemes being identified to make sure they are fed into the next decision process in the right order. Another example is the concept of ‘peakRate’ – the peaks in neural response associated with the onset-rhyme transitions in syllables (Oganian & Chang, 2019). Electrode sites where there is an enhanced peakRate response are conceptualised as labelling informative points in the speech stream where this information is used elsewhere in the system to focus processing attention on these transitions.

In a fully distributed recurrent process, as envisaged in a DisCo-like environment, none of these additional complexities arise. There is no computation of intermediate level cues which must then be transmitted to a higher-order process for interpretation and processing action. The informational value of the onset-rhyme transition, for example, is not something that needs to be signalled to some other process. The increase in neural activity at these points of transition is itself the increased computational weight the network assigns to these points in the speech signal. Where the phoneme order issue is concerned, the phonemically unsegmented articulatory dynamics that drive continuous speech interpretation mean that the phonological properties of the input are computed as the speech is heard. There is no re-encoding of the speech input as strings of phonemic labels and no stacking up of these labels until they can be interpreted – on this account, phonemic labels simply do not exist as neurocomputational entities that play a direct role in immediate speech interpretation.

### Conclusions

To answer the foundational questions we set out to address, we conclude with three hypotheses about the functional and neurobiological architecture that underpins the early processes of speech interpretation and about the computational vocabulary in terms of which these processes are conducted:

First, that the primary output of this process is an integrative mesoscale representation of the speech input that combines population-coded phonological and lexicosemantic constraints into an interpreted perceptual representation of the speech being heard.

Second, that the process of mapping from the incoming speech stream onto this interpreted representation is not mediated by a stratified hierarchy of intermediate representational transitions but by a non-hierarchical distributed neural architecture, where this architecture can be modelled, to a first approximation, in recurrent computational terms.

Third, and intrinsic to a recurrent architecture, that the distributional environment in which this interpretative mapping is learned - the statistical relationship of different phonological structures to different lexicosemantic outcomes – is encoded into the basic neural fabric of early speech analysis.

This combination of properties provides the basis, we suggest, for the earliness of speech interpretation – for the immediacy with which the listener hears what is being said.

## Acknowledgments

This research was funded by a European Research Council Advanced Investigator Grant to LKT under the European Community’s Horizon 2020 Research and Innovation Programme (2014–2022 ERC Grant Agreement 669820), and funded in whole, or in part, by the Wellcome Trust (Grant number 211200/Z/18/Z) to AC. We thank Ece Kocagoncu who collected the MEG data analysed here, Gareth Gaskell for his creative realisation of the original Distributed Cohort (DisCo) concept, Billi Randall, and the late Jeff Elman for advice and inspiration. For the purpose of open access, the authors have applied a CC BY public copyright licence to any Author Accepted Manuscript version arising from this submission.

## Notes

### Competing Interest Statement

The authors have declared no competing interest.

### Summary of Updates

Updated figures and revised interpretations

